# Examining Interspecific density-dependent dispersal in forest small mammals

**DOI:** 10.1101/2020.05.21.109827

**Authors:** Simon T. Denomme-Brown, Karl Cottenie, J. Bruce Falls, E. Ann Falls, Ronald J. Brooks, Andrew G. McAdam

## Abstract

The effects of conspecific densities on dispersal have been well documented. However, while positive and negative density-dependent dispersal based on conspecific densities are often shown to be the result of intraspecific competition or facilitation respectively, the effects of heterospecific densities on dispersal are examined far less frequently. This despite the potential for the analogous processes of interspecific competition and heterospecific attraction to influence dispersal. Here we use 51 years of live-trapping data on deer mouse (*Peromsycus maniculatus*), eastern chipmunk (*Tamias striatus*), red-backed vole (*Myodes gapperi*), and jumping mice (*Napaeozapus insignis* and *Zapus hudsonius*) to examine the effects of both conspecific and heterospecific densities on dispersal frequency. In terms of conspecific densities, jumping mice were more likely to disperse from areas of low conspecific densities, while red-backed voles and chipmunks did not respond to conspecific densities in their dispersal frequencies. When considering interspecific density effects, while there were no statistically clear effects of density on dispersal frequency, the effects of chipmunk and jumping mice densities on dispersal by red-backed vole were greater than the effects of conspecific densities, with voles more likely to disperse from areas of high chipmunk densities and low jumping mice densities. Likewise, the effect of chipmunk densities on dispersal by jumping mice was greater than the conspecific density effect. Conspecific densities clearly can affect dispersal by two of these four species, but the effects of heterospecific densities on dispersal frequency are less clear. Based on effect sizes it appears that there is potential for heterospecific effects on dispersal by some species in the community, but future experimental work could elucidate the strength and causes of these relationships.

## Introduction

Dispersal generally refers to either the movement of an individual from a place of origin to one of reproduction or from one reproductive locale to another (Greenwood 1980). This movement can influence fundamental ecological processes including population growth rates and viability, genetic connectivity and range expansions, and shapes spatial population dynamics (Greenwood 1980, Nathan 2006, Holyoak et al. 2008). However, the very nature of dispersal means it can be exceedingly difficult to examine in natural, empirical systems (Koenig et al. 1996), and thus there remain unanswered, fundamental questions in ecology regarding what drives organisms’ dispersal decisions (Sutherland et al. 2013).

The ultimate causes of dispersal are varied, as dispersal decisions are driven by both internal and environmental factors (Howard 1960). Dispersal is often thought to occur due to kin interactions or to avoid inbreeding. Additionally, individuals may disperse to improve living and breeding conditions when habitat quality varies either due to differences in the populations or communities existing in a patch, or due to variation in the intrinsic physical traits of habitat patches (Bowler and Benton 2005). One of the most oft-studied drivers of dispersal is variation in the density of conspecifics (Porter and Dooley 1993, Matthysen 2005). During emigration or settlement, patches of high conspecific densities may represent areas of potentially high intraspecific competition (Stamps 1991), leading individuals to emigrate from or avoid settling in such patches. This can create a pattern of positive density-dependent dispersal (DDD), where individuals benefit from reduced intraspecific competition and agonistic interactions when the conspecific densities at pre-dispersal sites are greater than those in post-dispersal settlement sites (Gaines and McClenaghan 1980, Porter and Dooley 1993, Matthysen 2005). This is the prevailing pattern observed in birds and mammals (Matthysen 2005). Alternatively, for species that exhibit facilitation in groups or that rely on access to resources that are scarce within a landscape the opposite can be true, with individuals dispersing more readily from areas of low density, termed negative DDD (Matthysen, 2005, Bowler and Benton 2005).

While dispersal can afford many benefits, it is not without its costs. Dispersers experience increased energy expenditure and physical wear, along with greater environmental risks including increased predation, habitat uncertainty and decreased resource security (Bonte et al. 2012). These costs can affect an individual at all three stages of a dispersal (i.e. emigration, transfer, and settlement). When faced with the risks associated with dispersal, it behooves an organism to be able to utilize any and all information from their surroundings as cues of possible fitness consequences of dispersal (Clobert et al. 2009). Such information can be acquired through an individual’s direct experiences with the abiotic environment, or in the form of social information (Danchin et al. 2004).

Social information is acquired through observation of how other individuals interact with a shared environment (Danchin et al. 2004). This information can be conveyed through direct signaling or, alternatively, through an individual’s performance in a given environment (Danchin et al. 2004). How others perform in an environment can convey public information regarding the quality of a habitat and associated resources, and there are numerous examples of organisms using the relative success of other individuals as indicators of their own fitness prospects (Danchin et al. 2004). For instance, many birds assess the suitability of habitat patches based on average reproductive success of conspecifics on a patch (i.e. habitat copying; Valone 1989, Doligez et al. 2002, Valone and Templeton 2002). The use of public information by dispersing individuals has the potential to markedly affect dispersal patterns, particularly when considered at the emigration and settlement stages of dispersal. For while variation in density of conspecifics can drive DDD through either competition or facilitation, the relative density and success of conspecifics may also provide public information to dispersers (Danchin et al. 2004). This public information can indicate high quality habitat or be conflated with high reproductive success leading to habitat copying (Danchin et al. 2004).

While variation in conspecific densities affects dispersal in numerous taxa (Matthysen 2005), the question of whether variability in the densities of heterospecifics can do the same is examined less often (Bowler and Benton 2005). Interspecific interactions can take a number of forms, including predation, competition and mutualisms, and provide the foundation for many ecological processes (Agrawal et al. 2007). Despite often being overlooked when examining possible causes of dispersal, there is evidence that interspecific interactions do have the potential to influence dispersal.

To date, examinations of interspecific interactions affecting dispersal have been largely restricted to predator-prey and parasite-host relationships (Bowler and Benton 2005, Chaianunporn and Hovestadt 2012). Examples include predator presence inducing the production of dispersing morphs in pea aphids (*Acyrthosiphon pisum*) (Sloggett and Weisser, 2002), and parasite infestation affecting recruitment and dispersal distances of great tits (*Parus major)* (Heeb et al. 1999). Additionally, Hauzy et al. (2007) demonstrated that dispersal by both prey and predator protist species were affected by the population density of the other. Notwithstanding these examples, literature on this topic remains sparse relative to investigations on conspecific densities. In particular, reviews of dispersal strategies have found few studies that focus on how interactions occurring within trophic levels, such as competition, might affect dispersal (Bowler and Benton 2005).

This dearth of literature is striking given that the presence of heterospecifics, particularly those sharing an ecological guild, is known to supply important information to many species regarding their environment (Mönkkönen et al. 1999, Seppänen et al. 2007, Cayuela et al. 2018). For instance, high densities of heterospecifics could indicate areas of increased interspecific competition and the possibility for positive interspecific DDD (Tilman 1987, Seppänen et al. 2007). Alternatively, in much the same way that habitat copying occurs amongst conspecifics, heterospecific attraction (Mönkönon et al. 1999) is exhibited by numerous species (Seppänen et al. 2007, Parejo et al. 2008, Cassaing et al. 2013, Szymkowiak et al. 2017), with increased density of heterospecifics causing animals to be less likely to disperse from or more likely to settle in an area. Heterospecific attraction occurs because heterospecific presence can help individuals assess the fitness prospects in possible breeding patches (Parejo et al. 2004) or indicate information about food resources (Seppänen et al. 2007). This process could feasibly lead to negative interspecific DDD, particularly because information provided by heterospecifics is often readily available as heterospecifics are often more plentiful in a landscape than conspecifics (Seppänen et al. 2007).

Examinations of the effects of heterospecific density on dispersal among members of the same trophic level have received relatively little examination to this point. Most studies of this type have been restricted to mesocosm experiments involving small invertebrates. De Meester et al. (2015) found that one of four species of nematodes involved in a mesocosm experiment dispersed earlier when interspecific population densities were high. However, a recent study of great crested newts (*Triturus cristatus*) has shown that newts disperse towards areas dense in heterospecifics (Cayuela et al. 2018), representing an example of heterospecific attraction affecting dispersal rates in wild vertebrate. Vertebrate studies on heterospecific attraction have otherwise been limited to its effect on habitat selection rather than dispersal (Parejo et al. 2008, Szymkowiak et al. 2017). Despite this, Cayuela et al.’s (2018) findings suggest that variation in heterospecific densities can indeed influence dispersal behaviours, but these effects have not been examined in other species or systems.

Here we used 51 years of trapping data to examine interspecific density effects on dispersal in four species of woodland rodents in a natural, temperate woodland. The most abundant species in this system has previously been shown to exhibit DDD based on conspecific densities (Denomme-Brown et al. *in review*). Additionally, while not as well established as in other taxa, there is evidence of heterospecific attraction in small mammals (Cassaing et al. 2013). We first examined the effects of conspecific densities on each species’ dispersal behaviours, and subsequently examined whether heterospecific population densities had similar or differing effects. This work represents one of examinations of intratrophic heterospecific DDD in a natural system (See also Cayuela et al. 2018) and the first to be performed on mammals. First, we expected dispersal frequency to be affected by density of conspecifics. Given the potential for heterospecifics to provide social information on habitat quality, we also expected that dispersal would be affected by density of heterospecifics. However, the value of conspecific versus heterospecific information is likely to depend on the abundance of a particular species. For a dominant species in a community, most information is likely to come from conspecifics, whereas for rare species more information is likely to be gained from heterospecifics (Sepännen et al. 2007). We, therefore, expected that the effect of heterospecific density would depend on the relative abundance of species in the community.

## Materials and Methods

### Study Area

Algonquin Provincial Park (APP) is a 765,300 ha provincial park located in central Ontario, Canada (45°35’03”N 78°21’30”W). From 1960-2015, small mammals were trapped in APP each year, with a variety of species caught regularly in the system. The analyses that follow in this paper will pertain to the four most abundant groups: the deer mouse (*Peromsycus maniculatus*), eastern chipmunk (*Tamias striatus*), red-backed vole (*Myodes gapperi*), and two species of jumping mice (*Napaeozapus insignis* and *Zapus hudsonius*). Other species caught in this system include red squirrel (*Tamiasciurus hudsonicus*), southern and northern flying squirrel (*Glaucomys volans* and *Glaucomys sabrinus*), meadow vole (*Microtus pennsylvanicus*), northern short-tailed shrew *(Blarina brevicauda*) and a number of shrew spp. in the genus *Sorex*.

### Data Collection

Trapping occurred on three consecutive nights either once or twice a month from May through September, up to a maximum of 10 three-night trapping sessions per year. Trapping initially occurred on 10 traplines, but an additional seven traplines were used beginning in 1995 such that the same 17 lines were trapped from 1995-2015. Distances between traplines varied, ranging from 115m – 16.9km apart.

Traplines consisted of 10 trapping stations set every 10m along transects. The study began with most lines having one trap per station, but by 1979 all lines had two traps per station, for a total of 20 traps per line, in order to avoid trap saturation. Traps were solely Sherman-style live traps (H.B. Sherman Traps Inc., Tallahassee, Florida) until 2013, when one Sherman trap at each station was replaced with a Longworth live trap (Natural Heritage Book Society, Totnes, Devon). Bait consisted of peanut butter and rolled oats until 1991, water-soaked sunflower seeds from 1991-2012 and water-soaked sunflower seeds in all traps and freeze-killed mealworms (*Tenebrio molitor*) in half the traps from 2013-2015. Mealworms and Longworth traps were used to decrease mortality in shrews (Do et al. 2013, Shonfield et al. 2013). Traps were set at dusk and checked after dawn. After being removed from a trap, individual mammals were weighed, sexed, aged and assessed for reproductive condition. They were then fitted with coded ear tags for individual identification upon recapture. Trapping protocols followed ASM guidelines (Sikes 2016) and were approved by the University of Guelph Animal Care Committee.

### Dispersal Detection

To be considered a dispersing individual within our analyses an animal must have been captured on a single trapline on at least two consecutive occasions, preceded or followed by at least one capture on a different trapline in the same trapping season. Such an individual was assumed to have been trapped while moving towards a trapline where it subsequently settled, or while dispersing from a trapline from which it either continued on or died. Thus, instances where an individual returned to its initial line of capture after moving between lines were not considered to be dispersal events. Individuals must have been captured a minimum of three times within a year in order to meet these criteria; all model analyses and calculations of dispersal frequencies included in this work were therefore performed using a dataset of individuals captured a minimum of three times. It should be noted that some loss of data led to the exclusion of five years from the dataset (1987-1991).

The straight-line distance between the first trapping station on each line was used to determine the distance between lines. While the strict criteria for being considered a dispersal event described in the preceding paragraph aimed to reduce errors, for additional quality control we also inspected the full capture history of all dispersing individuals for inconsistencies in other traits (e.g. sex and weight). We eliminated animals that appeared to disperse but that also exhibited unexplainable inconsistencies of this kind.

### Density measures

We calculated three measures of density for each species and each was converted from raw capture totals to captures per hundred trap-nights to account for variation in trapping effort both between years and among traplines. The first measurement was local trapline density, measured as the number of captures per hundred trap-nights at each trapline in each year. The second density measure was an regional trapline density, calculated annually as the mean number of captures per hundred trap-nights across all traplines in APP in that year.

Finally, we calculated the relative local trapline density for each trapline in each year by subtracting the regional density in a year from the local density on each line that same year. This relative local density measure was calculated to highlight the additional effects of local density, independent of annual variation in regional density, such as that caused by seed masting events that occur in this system (Falls et al. 2007). This relative measure of local density is also uncorrelated with regional density, allowing both the regional and relative local density measures to be used within the same models (*sensu* van de Pol and Wright 2009). Population densities were measured at both the local and regional scales because previous work in this system has shown that dispersal by deer mice is affected by an interaction between the densities existing at these two scales (Denomme-Brown et al. *in review*).

### Effects of Conspecific Densities on Dispersal Analysis

We assessed how each species’ propensity to disperse was affected by changes in conspecific densities by fitting sets of Generalized Linear Mixed Effects models (GLMM, binomial family, logit link) for each species. This method followed that performed in Denomme-Brown et al. (*in review*), and similarly, the response variable for all models was binary indicating whether each individual dispersed between lines or not. All models included the fixed effect of distance between the trapline where the animal was initially captured and the next closest trapline in the study system. This was included to account for the non-uniform spatial distribution of traplines and whether differences in movement among lines were primarily due to spatial effects on the probability of detecting a dispersal event (i.e. dispersal is more likely between nearby traplines). For the analysis of dispersal by red-backed voles and chipmunks we used the natural log of the raw distance in metres to partially account for non-linearities in the relationship between this variable and the probability of dispersal. For jumping mice, the raw distance in kilometres (km) was used. All models also contained the same random effects, with the initial trapline of capture, and the year of capture included to account for spatial and temporal variation in dispersal propensity.

In addition to the fixed and random affects already discussed, models had fixed effect structures composed of various combinations of sex, local density, relative local density and regional density. Some models also included interactions between local and regional densities or between local density and sex. For the analysis of red-backed voles, a significant non-linearity was identified between local density and the probability of dispersing. This non-linearity was accounted for by fitting a model with a quadratic term for local density. In all models in which they appeared, local density and regional density were converted to z-scores for each species (z = x − *x̄*/σ) based on the mean and standard deviation of all observations of conspecifics across years and lines. For details on the structure of all candidate models within this analysis see Supplementary Data SD1.

After fitting these sets of GLMMS, we used the Akaike Information Criterion (AIC) to assess the relative support for the candidate models fitted for each species (Burnham and Anderson 2007) by calculating AIC values for each model and then using ΔAIC values to perform model comparisons. Likelihood ratio tests (LRT) were used to assess the significance of the *year* and *trapline* random effects in each of the 8 candidate models. All models were assessed for overdispersion and collinearity of variables and were found to meet model assumptions. All GLMMs, in this section and those that follow, were fitted using R (Version 3.5; R Core Team, 2019), including the glmer function based in the lme4 package (Bates et al. 2015). For details on AIC comparisons between conspecific models see Supplementary Data SD2.

### Effect of Heterospecific Densities on Dispersal Analysis

Before assessing what the most appropriate combination of conspecific and heterospecific density effects were for best predicting the dispersal probabilities of each species, we utilized a simplified, consistent model structure to compare how strongly heterospecific densities might affect dispersal by each of the four small mammal species. This allowed us to make comparisons regarding the relative strength with which each species affected dispersal by the other three. For this analysis we used four GLMMs. The response variable for each of the four models was again a binary of whether or not an individual of a species dispersed, with each of the four models examining dispersal by one of deer mice, eastern chipmunk, red-backed voles or jumping mice. The fixed effects within each model were local conspecific density, and one term each for the local densities of each of the three remaining heterospecific species groups. As well, distance to nearest trapline was included as a fixed effect in each model. The random effects of year and initial trapline of capture were also included. For all models in the heterospecific analysis section of this study, model diagnostics and assumptions were checked in the same manner as in the intraspecific density-dependent models. Significance of random effects was determined using LRTs.

After comparing the relative strengths of heterospecific density effects on dispersal by the four species, we examined how best to predict the probability of dispersal for each species when considering heterospecific densities, while also considering the effects shown to be important in our conspecific analyses. To accomplish this, we again used a series of GLMMs, beginning with the best model for conspecific DDD by each species taken from our conspecific density analyses. The best model was determined via the lowest ΔAIC value. We used the model with the lowest ΔAIC in order to reduce the number of comparisons that would result from performing these analyses with the multiple base models for which AIC had shown strong support. We then constructed six candidate heterospecific models per species for this analysis. We used the best conspecific model as the first candidate model, and then created three additional candidate models by separately adding the fixed effect of raw local density of each of the non-focal species to the conspecific model. The fourth candidate model consisted of the best conspecific model for the focal species with the added fixed effect of cumulative local density of all three of these heterospecific populations combined. The final candidate model consisted of the best conspecific model for that species with the addition of an interaction between cumulative local density of the three heterospecific populations and sex. This interaction was included as male and female individuals might be expected to respond differently to heterospecific presence based on intersex differences in reproductive strategies. After the six models for each of the four species were fitted, we again performed AIC comparisons using ΔAIC values in order to determine the model with the best support. LRTs were again used to assess the significance of random effects. For details on AIC comparisons and model structure see Supplementary Data SD3.

## Results

### Population densities and frequency of dispersal

Summary statistics regarding deer mice are reported in Denomme-Brown et al. (*in review*, see Supplementary Data SD4). Of the possible dispersers for each species, 3.2% of eastern chipmunks (n = 40/1239), 2.7% of red-backed voles (n = 21/770) and 6.7% of jumping mice (n = 31/464) dispersed between traplines. The range of distances travelled by the small mammals varied widely. eastern chipmunks, red-backed voles and jumping mice exhibited dispersal distance ranges of 0.115 – 6.34km, 0.115 – 10.83km and 0.115 – 12.48km respectively (Figure 1). On average, red-backed voles moved the farthest (*x̄* ± *SE* = 1.27 ± 0.52 km), followed by chipmunks (*x̄* ± *SE* = 1.19 ± 0.28 km) and jumping mice (*x̄* ± *SE* = 1.01 ± 0.43 km,). As two lines within the system are particularly close to one another (115m), it is worth noting that 14.3% of red backed vole dispersals, 17.5% of chipmunk dispersals and 29% of jumping mice dispersals occurred between these two lines.

**Figure 1:**
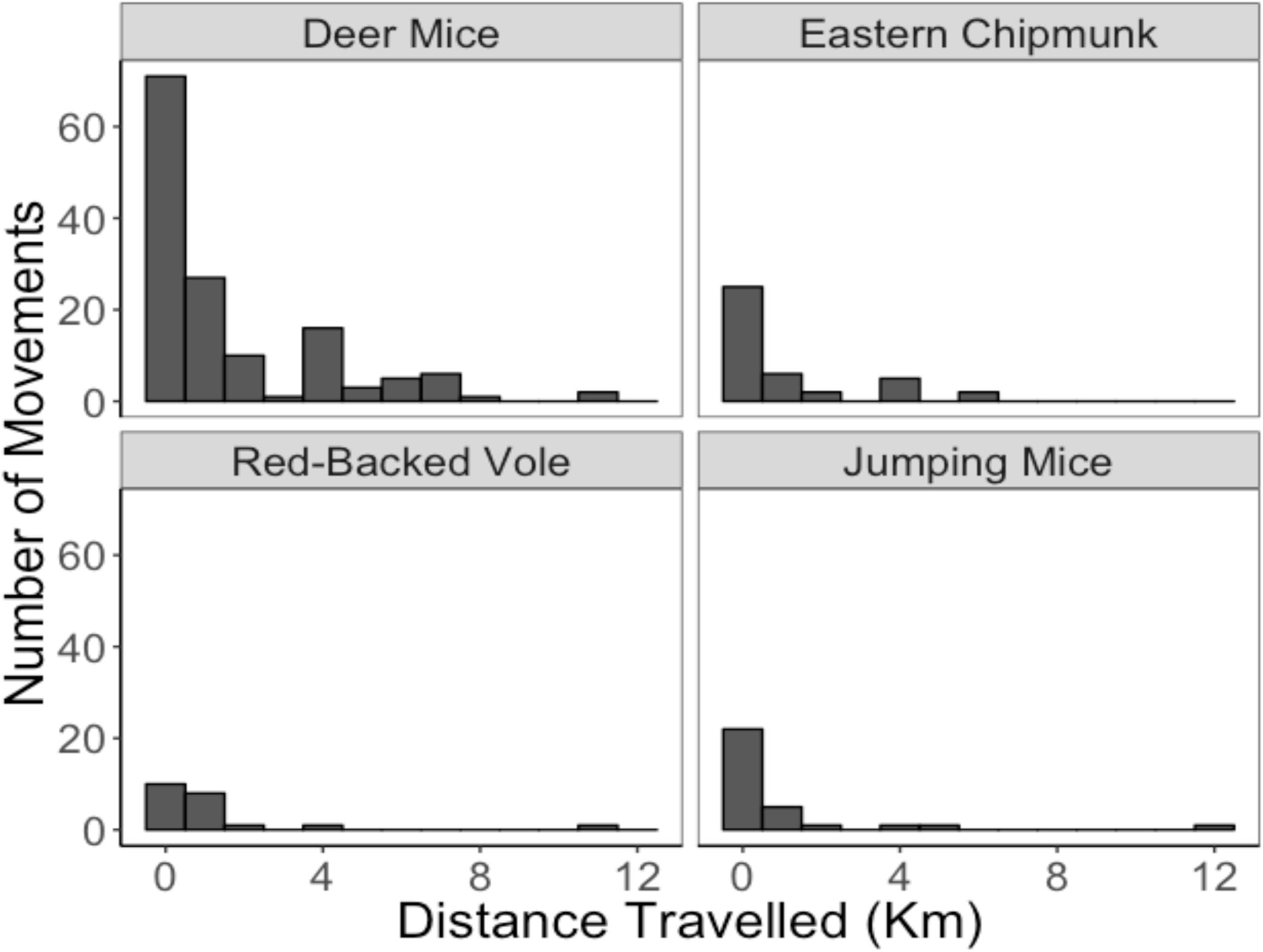
Dispersal distances of small mammals in APP from 1960-2015.

The densities of all these species in APP varied quite markedly through the time series (Figure 2). In general, deer mice were the most abundant (see Denomme-Brown et al. *in review*; Supplementary Data SD4). Beyond deer mice, eastern chipmunk were regionally most abundant on average and their abundances also varied most widely (*x̄* = 3.36 capt/100 trapnights, *SD* = 3.34, range = 0 −14.51). Red backed voles were next most abundant (*x̄* = 2.75 capt/100 trapnights, *SD* = 2.1, range = 0.11 − 8.01 capt/100 trapnights), followed by jumping mice (*x̄* = 2.24 capt/100 trapnights, *SD* = 2.12, range = 0 − 7.25 capt/100 trapnights) (Figure 2).

**Figure 2:**
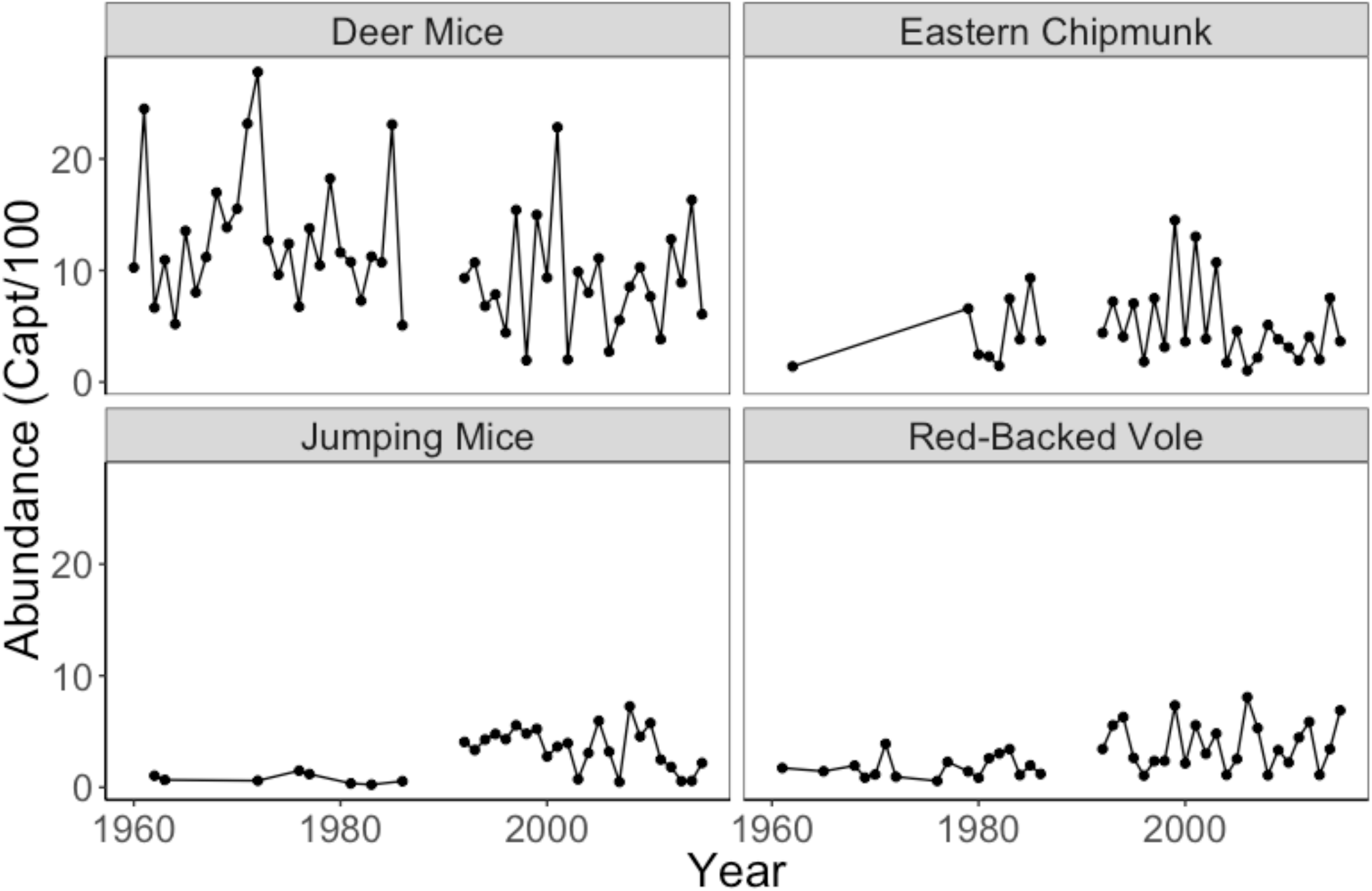
Abundance of small mammals in captures/100 trapnights in APP from 1960-2015. Vertical black lines signify five years where data were not available.

### Conspecific Dispersal Models

The best model for deer mouse dispersal based on AIC scores is fully reported in Denomme-Brown et al. (*in review*, fixed effects in Table 1). This model included significant negative effects of both local density and regional density on dispersal; with a positive interaction between local and regional density, indicating that the negative effect of local density on dispersal was more pronounced in years of low regional density.

**Table 1:** Fixed effects included in best conspecific model for density-dependent dispersal by deer mice (*Peromyscus maniculatus*), eastern chipmunk (*Tamias striatus*), red-backed vole (*Myodes gapperi*), and woodland and meadow jumping mice (*Napaeozapus insignis* and *Zapus hudsonius*) in Algonquin Provincial Park from 1960-2015. (+) indicates a positive effect while (−) represents a negative effect. (*) indicates the effect is significant at α = 0.95. NA indicates the fixed effect was not included in the model. (:) indicates an interaction between the two stated fixed effects.

| Species | Distance to closest line | Sex (male) | Local conspecific density | Regional conspecific density | Local conspecific density:regional conspecific density | local conspecific density:sex |
| --- | --- | --- | --- | --- | --- | --- |
| deer mice<sup>†</sup> | —* | +* | —* | —* | +* | +* |
| eastern chipmunk | —* | + | NA | NA | NA | NA |
| red-backed vole | — | NA | NA | NA | NA | NA |
| jumping mice | —* | +* | —* | NA | NA | NA |

<sup>†</sup> model presented in Denomme-Brown et al. *in review*

The conspecific model for chipmunk dispersal with the lowest AIC score had sex of individual as a fixed effect, in addition to a term for the distance between the initial trapline of capture and the next nearest the trapline in the study. The probability of dispersal decreased with increasing distance to the nearest trapline as would be expected (*β* ± *SE* = −3.4 ± 0.64, *Z* = −5.3, *P* < 0.0001), however, sex (*β* ± *SE* = 0.51 ± 0.33, *Z* = 1.53, *P* = 0.13) was not a significant predictors of dispersal. The random effects of *year* and *trapline* were both not significantly different from zero (LRT: *year*: Deviance = 319, *X^2^_1_* = 0, *P* = 1; *trapline*: Deviance = 319, *X^2^_1_* = 0, *P* = 1). This model explained 9.96% of the overall deviance. There was also strong support based on AIC values (ΔAIC < 2) for four other models, but no density terms were significant in any of these models, nor did the direction or significance of the fixed effects change from those in the best model change.

The best model for red-backed vole dispersal based on AIC scores included only the fixed effect of distance to closest trapline. The probability of dispersal by red-backed voles decreased with increasing distance to nearest trapline, but this effect was not significant (*β* ± *SE* = −0.13 ± 0.52, *Z* = −0.28, *P* = 0.78). The random effects of *year* and *trapline* were both not significantly different from zero (LRT: *year*: Deviance = 216, *X^2^_1_* = 0, *P* = 1; *trapline*: Deviance = 215, *X^2^_1_* = 0, *P* = 0.25). Overall, this model explained 1.2% of the deviance. There was also strong support based on AIC values (ΔAIC < 2) for three additional models. The first only contained the nonsignificant fixed effects of sex of the individual (*β* ± *SE* = −0.54 ± 0.46, *Z* = 1.16, *P* = 0.25) and distance to closest trapline (*β* ± *SE* = −0.14 ± 0.54, *Z* = −0.26, *P* = 0.80). The second again contained nonsignificant fixed effects of sex of the individual (*β* ± *SE* = −0.57 ± 0.47, *Z* = 1.23, *P* = 0.22), and distance to closest trapline (*β* ± *SE* = −0.17 ± 0.49, *Z* = 0.34, *P* = 0.73), along with the nonsignificant quadratic effect of relative local density (Linear: *β* ± *SE* = − 0.37 ± 0.27, *Z* = −1.35, *P* = 0.18; Quadratic *β* ± *SE* = 0.32 ± 0.17, *Z* = 1.94, *P* = 0.052). In the third model with strong support, a significant positive interaction (*β* ± *SE* = 0.11 ± 0.05, *Z* = 2.08, *P* = 0.038) between the negative effects of both relative local density (*β* ± *SE* = −0.10 ± 0.05, *Z* = −1.74, *P* = 0.08) and regional density (*β* ± *SE* = −0.38 ± 0.37, *Z* = −1.03, *P* = 0.30), suggested that in years of low regional density voles were more likely to disperse from areas of low relative local density, and that in years of high regional density, voles were more likely to disperse from areas of high relative local density (Figure 3). The effects of sex and distance to closest trapline were again both not significant (Sex: *β* ± *SE* = 0.66 ± 0.61, *Z* = 1.28, *P* = 0.20; Closest Trapline: *β* ± *SE* = −0.16 ± 0.44, *Z* = −0.36, *P* = 0.71).

**Figure 3:**
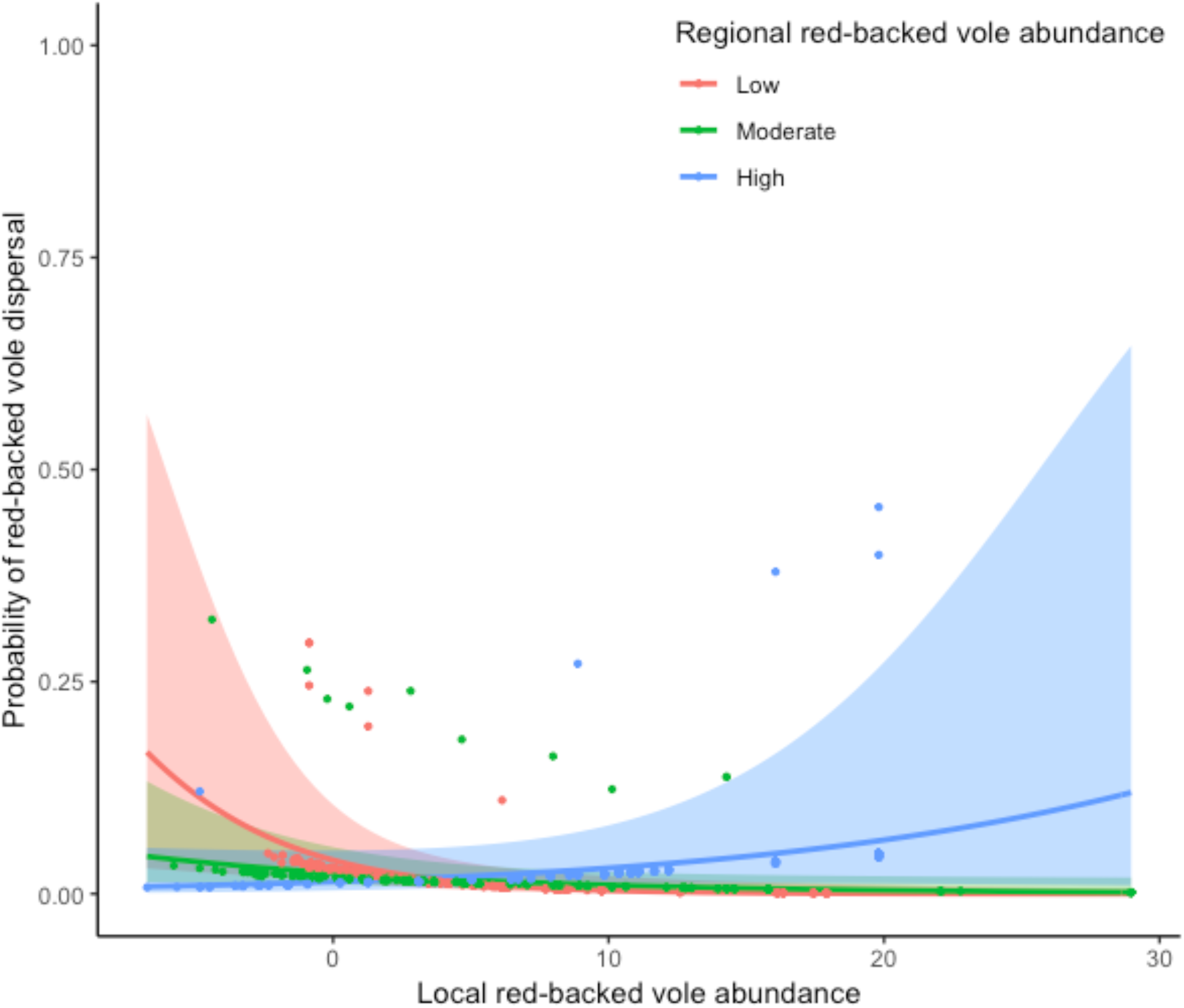
Effect of the interaction between relative local red-backed vole (*Myodes gapperi)* density and regional red-backed vole density on dispersal by red-backed vole (*β* ± *SE* = 0.11 ± 0.05, *Z* = 2.08, *P* = 0.038) in a conspecific model with strong support based on AIC values. Voles appear more likely to disperse from areas of low relative local density in years of low regional density; and from areas of high relative local density in years of high regional density. Abundances scaled from captures/100 trapnights. Shading represents 95% CI..

For jumping mice, the best model included the fixed effects of local density, sex of individual and distance to nearest trapline. Jumping mice were more likely to disperse when local densities were low (*β* ± *SE* = −0.62 ± 0.24, *Z* = −2.53, *P* = 0.011, Figure 4A), and if they were male (*β* ± *SE* = 0.89 ± 0.40, *Z* = 2.24, *P* = 0.025). As in the chipmunk model, the probability of dispersing significantly decreased with increasing distance to the next closest trapline (*β* ± *SE* = −3.03 ± 0.87, *Z* = −3.50, *P* < 0.001). The random effects of *year* and *trapline* were both not significantly different from zero (LRT: *year*: Deviance = 200, *X^2^_1_* = 0, *P* = 1; *trapline*: Deviance = 200, *X^2^_1_* = 0.04, *P* = 0.84). This model explained 12.15% of the overall deviance. There was also strong support based on AIC values (ΔAIC < 2) for two other models. In the first, like in the model reported above, males were significantly more likely to disperse (*β* ± *SE* = 0.91 ± 0.40, *Z* = 2.31, *P* = 0.02), and the probability of dispersal decreased with increased distance to the nearest trapline (*β* ± *SE* = −2.95 ± 0.90, *Z* = −3.29, *P* = 0.001). Unlike the model with the lowest AIC score, this model included relative local density rather than local density and, similarly, dispersal probability decreased with relative local density (*β* ± *SE* = −0.16 ± 0.07, *Z* = −2.43, *P* = 0.015). The other model with strong support found that probability of dispersal decreased was negatively associated with relative local density (*β* ± *SE* = −0.16 ± 0.07, *Z* = −2.38, *P* = 0.018) and distance to closest trapline (*β* ± *SE* = −3.0 ± 0.89, *Z* = −3.38, *P* < 0.001), was more likely in males (*β* ± *SE* = 0.90 ± 0.40, *Z* = 2.27, *P* = 0.023), and was not significantly affected by regional density (*β* ± *SE* = −0.15 ± 0.19, *Z* = −0.78, *P* = 0.43).

**Figure 4:**
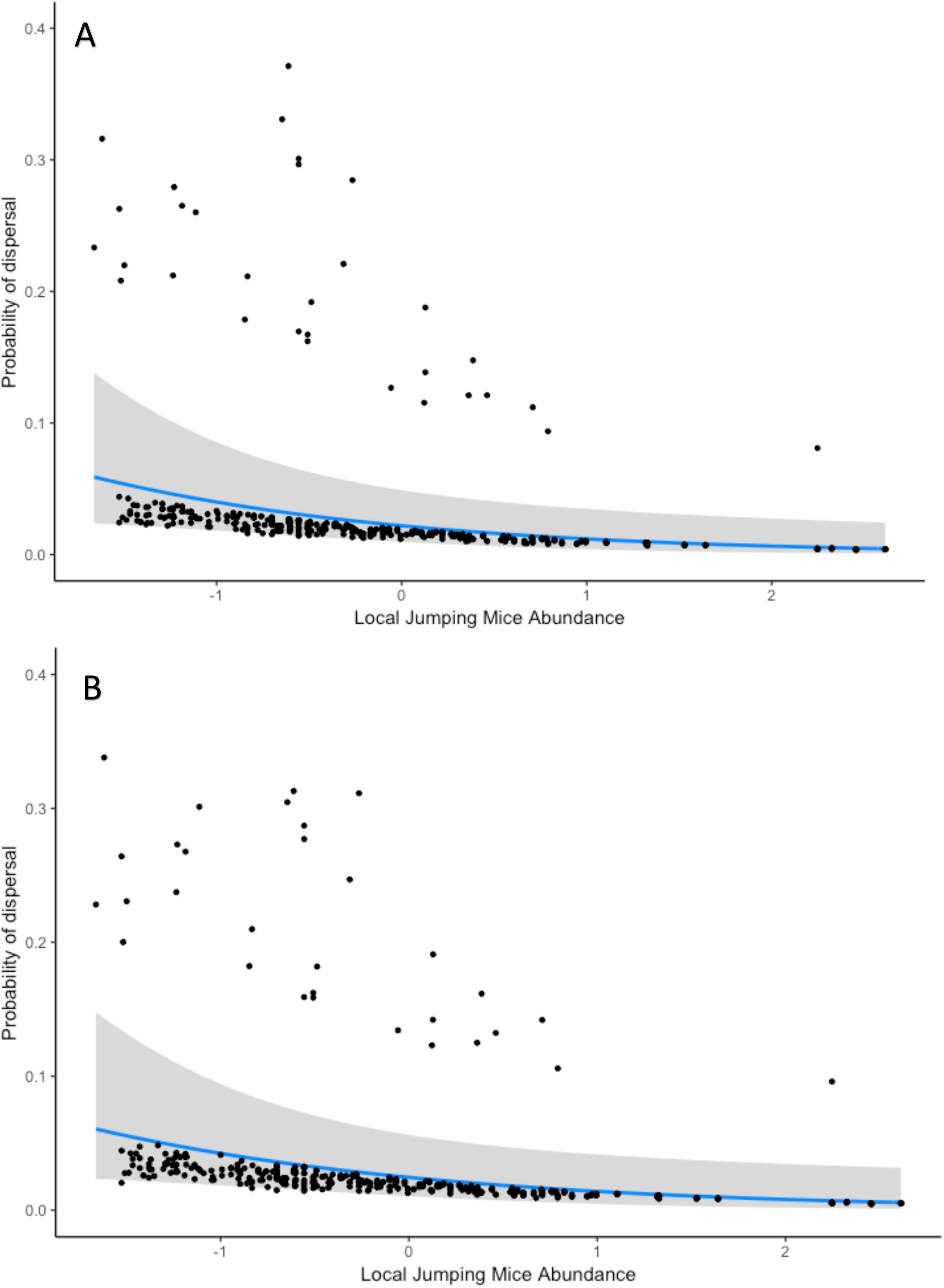
A) Jumping mice (*Napaeozapus insignis* and *Zapus hudsonius*) were more likely to disperse at low local densities (*β* ± *SE* = −0.62 ± 0.25, *Z* = −2.53, *P* = 0.012; conspecific model) B) Jumping mice were again more likely to disperse at low conspecific densities in the heterospecific model (*β* ± *SE* = −0.57 ± 0.25, *Z* = −2.28, *P* = 0.023). Abundances scaled from captures/100 trapnights. Shading represents 95% CI.

### Heterospecific dispersal models

The four consistently structured heterospecific density models used to compare the strength of heterospecific density effects on dispersal by the four mammal species groups indicated varying importance of heterospecific effects on dispersal. Effect sizes and associated standard errors are presented in Table 2, along with indications of significance. In general, deer mice dispersal was most strongly affected by conspecific density (*β* ± *SE* = −0.50 ± 0.13, *Z* = −3.89, *P* = 0.0001); chipmunk dispersal was most strongly affected by jumping mice densities (*β* ± *SE* = −0.18 ± 0.18, *Z* = −0.98, *P* = 0.32) and deer mice densities (*β* ± *SE* = −0.14 ± 0.18, *Z* = −0.78, *P* = 0.43), although these effects were not significant. Red-backed vole dispersal was most strongly affected by jumping mice densities (*β* ± *SE* = −0.47 ± 0.32, *Z* = −1.45, *P* = 0.15) and chipmunk densities (*β* ± *SE* = 0.32 ± 0.24, *Z* = 1.34, *P* = 0.18), although neither effect was significant; jumping mice dispersal was most strongly affected by chipmunk densities (*β* ± SE = −0.94 ± 0.49, *Z* = −1.93, *P* = 0.054), followed by conspecific densities (*β* ± *SE* = −0.62 ± 0.27, *Z* = −2.32, *P* = 0.020). For full model results see Supplementary Data SD5.

**Table 2:** Effect of both conspecific and heterospecific local density on dispersal by deer mice (*Peromyscus maniculatus*, DM), eastern chipmunk (*Tamias striatus*, CHIP), red-backed vole (*Myodes gapperi*, RBV), and woodland and meadow jumping mice (*Napaeozapus insignis* and *Zapus hudsonius*, JM) in Algonquin Provincial Park from 1960-2015. (*) indicates the effect is significant at α = 0.95. (†) indicates that the *P* value for this effect is 0.05.

|  |  | Fixed Effects |  |  |  |
| --- | --- | --- | --- | --- | --- |
|  |  | DM | CHIP | RBV | JM |
| Focal<br>Dispersers | DM | $-0.50 \pm 0.13^*$ | $-0.099 \pm 0.14$ | $0.045 \pm 0.084$ | $0.052 \pm 0.10$ |
| | CHIP | $-0.14 \pm 0.18$ | $-0.029 \pm 0.23$ | $0.015 \pm 0.14$ | $-0.18 \pm 0.18$ |
| | RBV | $-0.11 \pm 0.28$ | $0.32 \pm 0.24$ | $-0.16 \pm 0.29$ | $-0.47 \pm 0.32$ |
| | JM | $0.12 \pm 0.25$ | $-0.94 \pm 0.49^\dagger$ | $-0.17 \pm 0.23$ | $-0.62 \pm 0.27^*$ |

When examining how to best predict dispersal using both conspecific and heterospecific densities, the addition of terms for heterospecific densities did little to improve upon the best models for conspecific DDD in deer mice, chipmunks or red-backed voles. For all three of those species, after assessing the best conspecific model and the five models containing heterospecific density terms, the conspecific model remained the model with the lowest AIC score. Two other heterospecific models for deer mice dispersal, four others for chipmunks and two others for red-backed vole also had strong support based on ΔAIC values (ΔAIC < 2). However, none of the heterospecific density terms were significant in any of these models.

For jumping mice, the model with the lowest AIC score when assessing the effects of heterospecific densities on dispersal consisted of the best model from the conspecific jumping mouse analysis with the addition of the fixed effect of local chipmunk density. Like the model for red-backed voles, heterospecific population densities did not significantly affect probability of dispersal in this jumping mouse model. While jumping mouse dispersal decreased with increasing chipmunk abundance, this effect was not significant (*β* ± *SE* = −0.74 ± 0.44, *Z* = −1.70, *P* = 0.090, Figure 5). Like in the base conspecific model, sex was again a significant predictor of dispersal by jumping mice in this model (Sex (male) *β* ± *SE* = 0.84 ± 0.40, *Z* = 2.12, *P* = 0.034, Figure 4B). The significance and direction of the remaining fixed effects were the same as those described in the base conspecific model (Local Conspecific Density: *β* ± *SE* = −0.57 ± 0.25, *Z* = − 2.28, *P* = 0.023; Distance to nearest trapline: *β* ± *SE* = −2.23 ± 1.004, *Z* = −2.22, *P* = 0.027). The random effects of *year* and *trapline* were both not significantly different from zero (LRT: *year*: Deviance = 197, *X^2^_1_* = 0, *P* = 1; *trapline*: Deviance = 196, *X^2^_1_* = 0.56, *P* = 0.45). The model explained 13.82% of the deviance. The two models with the additional fixed effects of total local competitor density, and the interaction between local competitor density and sex respectively each had strong support based on ΔAIC (ΔAIC < 2). However, the heterospecific effects in these models were not significant. The model with no heterospecific fixed effects also had strong support.

**Figure 5:**
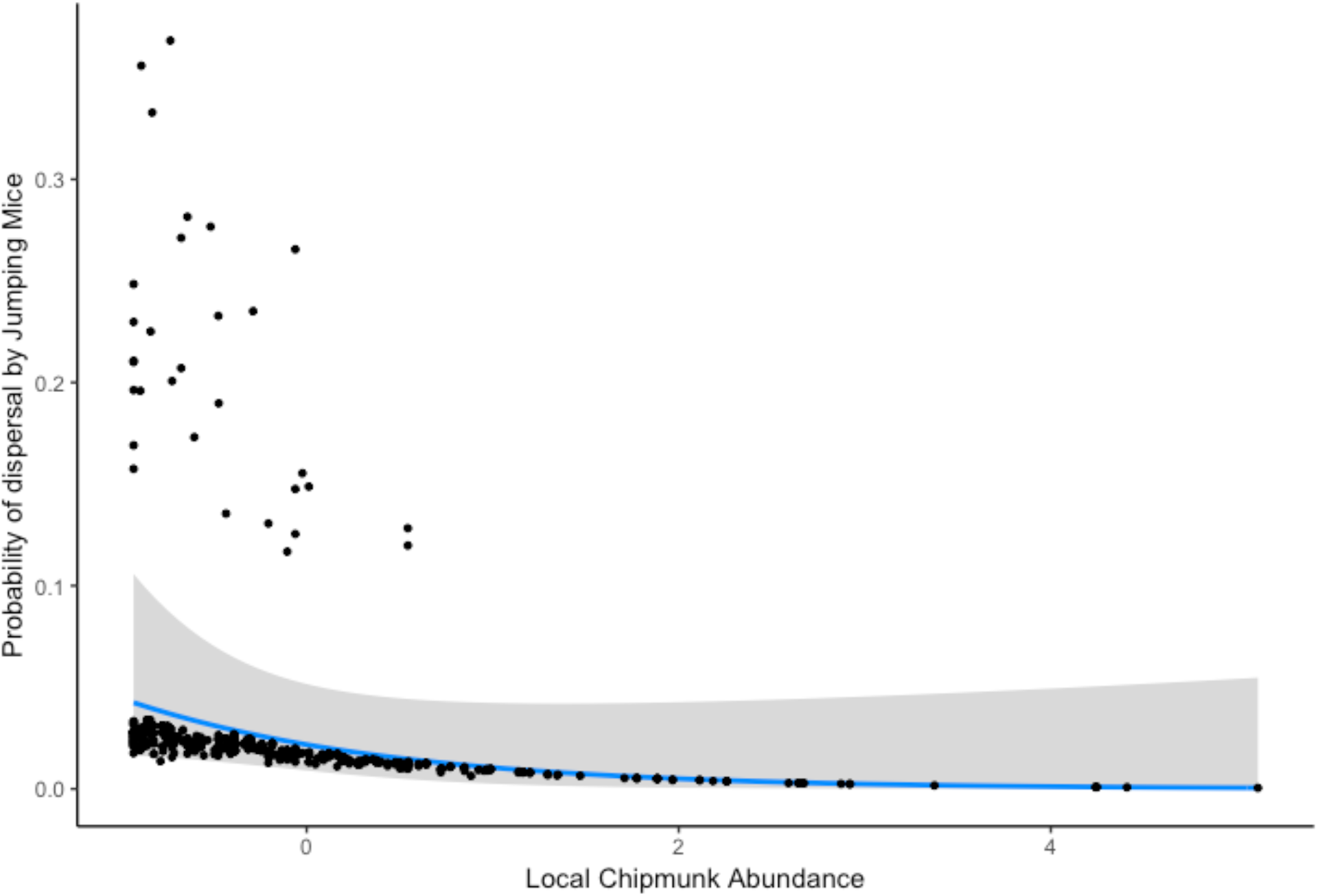
Effect of local eastern chipmunk (*Tamias striatus*) abundance on dispersal by jumping mice (*Napaeozapus insignis* and *Zapus hudsonius*) in APP between 1960-2015 (heterospecific model). The effect was not significant (*β* ± *SE* = −0.74 ± 0.44, *Z* = −1.70, *P* = 0.090) but did improve the model based on AIC. Abundances scaled from captures/100 trapnights. Shading represents 95% CI.

## Discussion

### Conspecific Density Effects on Dispersal

Overall, conspecific densities affected dispersal probability of jumping mice; chipmunks did not appear to alter their dispersal frequency based on variation in conspecific densities, and the effects of conspecific densities on dispersal by red-backed voles are unclear. These different responses suggest differences in these species’ natural history influence their response to density. As each of these three species exhibited marked changes in population density throughout our study (Figure 2), these different responses are unlikely to derive from inadequate variation throughout the time series.

Dispersal by jumping mice decreased with increased conspecific densities, an example of negative DDD. While positive DDD is more common in mammals (Matthysen 2005), there are numerous examples of negative DDD in mammals, and specifically in rodents (Denomme-Brown et al. *in review*, Rehmeier et al. 2004, Matthysen 2005). Numerous explanations for negative DDD have been suggested including attraction to conspecifics (Danielson and Gaines 1987, Stamps 1991) and exclusion of immigrants in periods of high density (i.e. “social fences”; Hestbeck 1982). Dispersal by jumping mice in general is very rarely studied (but see Bowman et al. 2001, Vignieri 2005, Vignieri 2007). Given this, additional work is needed to shed light on the exact cause of negative DDD in these rodents.

DDD by red-backed voles is less clear as the model with the lowest AIC score contained no density terms, but there was also strong support for the model containing a significant interaction between regional density and relative local density. In this model, the effect of relative local density on dispersal hinged on the regional density in the system at the time. It appears that voles may disperse more from areas of low relative conspecific density in years of low regional conspecific density, and more often from areas of high relative conspecific density in years of high regional conspecific density (Figure 3). Voles in APP could be responding to multiple drivers of DDD, perhaps dispersing at high density to avoid increased agonistic interactions (Gaines and McClenaghan 1980), and at low densities either to find mates when they become scarce or more suitable habitats (Stamps 1991). However, while this interaction is statistically significant, the effect size is quite small and given the very few dispersing individuals we observed for this species (n = 21) it appears that a few outliers may be driving this relationship (Figure 3).

Unlike the previous two rodents, the eastern chipmunk exhibited no significant change in dispersal frequency in response to varying conspecific densities. While red-backed voles and jumping mice maintain overlapping territories, chipmunks are highly territorial and aggressive in territory defence (Getty 1981, Lacher and Mares 1996). This highly territorial nature could perhaps make them more prone to exhibiting positive DDD, as increasingly frequent aggressive interactions due to increased density can lead to dispersal (Gaines and McClenaghan 1980, Porter and Dooley 1993, Matthysen 2005). There is to our knowledge no work done on how differing levels of territoriality might influence the direction or intensity of DDD. However, it could be that given their aggressive territorial defence, chipmunks are less affected by loss of resource access that might drive dispersal by other less territorial species under similar conditions, leading to them being less affected by density in terms of their dispersal, resulting in the nonsignificant relationship observed here.

### Heterospecific Density Effects on Dispersal

Based on our analyses, it appears that there are weak, albeit notable effects of heterospecific density on dispersal in some of the species examined. While the best conspecific model for jumping mice was improved by the inclusion of a heterospecific density term for red-backed voles, the effect of local red-backed vole density on dispersal by jumping mice did not represent a statistically clear relationship. This lack of evidence of heterospecific effects of DDD in these best predictive models may indicate that these species are not strongly affected by the density of heterospecifics when making dispersal decisions; however, an examination of the effect sizes of these best predictive models, as well as in the four consistently structured models of heterospecific dispersal affects, suggest that heterospecific densities may still play role in influencing dispersal.

In the case of the four consistently-structured heterospecific models, deer mice were the only species where the effect of conspecific densities on dispersal far outweighed the effects of heterospecifics (Table 2). The strongest effects on dispersal by chipmunks were the density of conspecifics and jumping mice, red-backed vole dispersal was more strongly affected by jumping mice and chipmunk densities than conspecific densities, and jumping mice dispersal was most affected by chipmunk density. It is also worth noting that chipmunks in particular had greater effects on dispersal by red-backed vole and jumping mice than they did on their own dispersal. Red-backed voles were more likely to disperse when chipmunks were in high abundance, perhaps indicating some level of interspecific competition, while jumping mice were more likely to disperse when chipmunks were at low abundances. In a similar fashion, jumping mice affected red-backed vole dispersal at the same effect size as they influenced their own dispersal. In the case of the best predictive model for jumping mice, it is notable that the effect size of chipmunk density on jumping mice dispersal was greater than the effect of conspecific density, despite not being a statistically clear relationship. While some of these effects do not meet the requirements for statistical significance, the improvement of models based on AIC scores and the substantial effect sizes of heterospecific densities when compared to conspecific densities in a consistently structured set of models suggest that heterospecific densities do have some effect on dispersal probabilities.

There are multiple possible explanations for why heterospecific densities might influence dispersal in these species. Perhaps most intuitively, resource competition might lead to increased density of heterospecifics causing increased dispersal (Tilman 1987, Seppänen et al. 2007). However, nearly all the effects of heterospecific density on dispersal in the consistently structured models are negative, with more numerous heterospecifics in the environment leading to reduced frequency of dispersal (Table 2). This reduced emigration in the face of increased competitor presence could stem from the use of heterospecific social information and subsequent habitat copying (Doligez et al. 2002, Valone and Templeton 2002, Danchin et al. 2004). Alternatively, it could be that high densities of competitors lead to reduced access to resources, resulting in individuals of poorer condition. In order to disperse, individuals require both the capacity and motivation to do so (Bowler and Benton 2005, Benard and McCauley 2008). Dispersal distance has been shown to be correlated with body size in mammals (Sutherland et al 2000), and reaching a threshold in body size is crucial for dispersal to occur in some rodents (Holekamp 1986). If increased density of competitors does negatively influence body size, then it could lead to reduced dispersal from areas where heterospecifics are plentiful. Further work examining individual level variation amongst dispersers and non-dispersers would help elucidate which mechanism might be driving these observations.

It should be noted that it is likely our study design, despite spanning an extensive time-series, may still not be ideal for detecting dispersal responses to heterospecific densities regardless of their presence. As previously discussed, numerous species use public information provided by heterospecifics in order to make behavioural decisions (Mönkkönen et al. 1999, Parejo et al. 2004, Seppänen 2007), including dispersal decisions (Cayuela et al. 2018). Information provided by heterospecifics is considered especially useful as heterospecifics are generally more abundant in an organism’s environment than conspecifics (Sepännen 2007). By tapping into public information, an organism is able to better assess and respond to its’ environment than if relying solely on information from conspecifics. This reasoning provides a clue as to why we may not have detected a strong signal in this study. Deer mice are by far the most abundant species in the system, and we subsequently have the most data regarding their dispersal in this system, including the highest number of dispersal events. While our data may be sufficient to detect heterospecific effects on dispersal by deer mice, as the most abundant species in the landscape they are less likely to rely on heterospecifics for information, when they can glean more precise information regarding habitat suitability from consistently abundant conspecifics (Sepännan 2007). This would leave the three other species as good candidates for examining heterospecific effects on dispersal. Unfortunately, it is likely that having significantly less data for these species compared to deer mice means that for these species, while the inclusion of heterospecific density terms may have large effect sizes and improve models based on AIC scores, we are unlikely to be able to detect heterospecific signals with any statistical significance using these data.

### Distance and Frequency of Dispersal

Our observations of dispersal in this study represent numerous record dispersal distances. Our maximum recorded distance moved by a red-backed vole (10.8 km) is greater than both the greatest homing (600m; Bovet 1980) and non-homing (494m; Bowman 2001) movements previously reported for this species. The greatest movement we observed by a jumping mouse (12.48km) was not identified to species, but exceeded the record movement previously reported by either of the two jumping mice species in this study (>800m by *N. insignis*, Ovaska and Herman 1988). The longest dispersal event by eastern chipmunk that we were able to identify in the literature was a homing movement of 0.55 km (Seidel 1961), and is far outstripped by our observation of a 6.33km dispersal. These record-breaking movements suggest that, like the movements by deer mice reported by Denomme-Brown et al (*in review*), the tail of the dispersal distance curve for many small mammal species has been grossly underestimated to this point.

The frequency of dispersal by species in our analyses was largely consistent with those presented in other studies. Jumping mice dispersed most frequently with 6.7% percent of individuals dispersing. This finding is not drastically different from that of Bowman et al. (2001) who found that 9.4 % woodland jumping mice in their trapping study undertook long-distance dispersals (>100m), which was more than either red-backed voles (1.8%) or deer mice (4.2%).We observed marginally more frequent movements by red backed voles (2.7%) compared to Bowman et al. (1.8%; 2001), although this difference is minimal and is consistent with this species dispersing the least frequently of the three discussed here so far. The frequency of movement events by chipmunks has seldom been reported in the literature outside of their interactions with roads (Ford and Fahrig 2008) or ranging for space use (Bowers 1995). As such, there are not estimates of dispersal frequencies that we could acquire with which to compare our frequencies of chipmunk dispersal.

In conclusion, most rodents examined in this study appear to respond to conspecific densities in terms of their dispersal. The responses in dispersal by these same species to variability in heterospecific densities was less clear, but for jumping mice the inclusion of heterospecific density terms improved the best model for dispersal probability based on AIC scores, and in a set of consistently-structured models dispersal by eastern chipmunks, red-backed voles and jumping mice were all affected by heterospecific densities as much or more than by conspecific densities. The causes of the different responses to conspecific density are difficult to discern given the lack of information on dispersal in these species. Work examining or summarizing how differing levels of territoriality affect how different species exhibit DDD would aid in answering why these species responded so differently to variation in conspecifics. As for the effects of heterospecific densities on dispersal, it appears that while the effects may be weak, heterospecific densities can influence dispersal probability in some species. Future work involving field experimentation that accounts for individual-level variation may be able to better elucidate the causes and strength of these relationships.

## Acknowledgements

We would like to acknowledge the countless individuals that worked to collect the data analyzed here over more than fifty years. Thank you to the staff at the Algonquin Wildlife Research station for providing logistical support throughout the study.

Funding: The field work in this study was funded in part through the support of the National Science and Engineering Research Council of Canada (NSERC) and the Ontario Ministry of Natural Resources. Funding for STDB is provided in part by an NSERC post graduate scholarship.

## Supplementary Data

SD1—conspecific density dependent dispersal candidate model structure

SD2—conspecific density dependent dispersal model AIC comparisons

SD3—heterospecific density dependent dispersal candidate model structure and output

SD4—summary statistics regarding deer mice in APP (taken from Denomme-Brown et al. in review)

SD5—Full model results for consistently structured heterospecific density-dependent dispersal models

## Literature Cited

Agrawal, A.A. et al. 2007. Filling key gaps in population and community ecology. Frontiers in Ecology and the Environment 5: 145–152

Bates, D., M. Mächler, B. Bolker, and S. Walker. 2015. Fitting linear mixed-effects models using lme4. Journal of Statistical Software 67:1–48.

Benard, M. F. and S. J. McCauley. Integrating across life-history stages: consequences of natal habitat effects on dispersal. The American Naturalist 171: 553–567.

Bonte, D. et al. 2012. Costs of dispersal. Biological Reviews 87:290–312.

Bovet, J. 1980. Homing behaviour and orientation in the red-backed vole, *Clethrionomys gapperi*. The Canadian Journal of Zoology 58:754–760.

Bowers, M.A. 1995. Use of space and habitats by the eastern chipmunk, *Tamias striatus*. Journal of Mammalogy 76:12–21.

Bowler, D. E., and T. G. Benton. 2005. Causes and consequences of animal dispersal strategies: relating individual behaviour to spatial dynamics. Biological reviews of the Cambridge Philosophical Society 80:205–225.

Bowman, J., G. J. Forbes, and T. G. Dilworth. 2001. Distances moved by small woodland rodents within large trapping grids. The Canadian Field Naturalist 115:64–67.

Burnham, K.P. and D. R. Anderson. 2007. Model selection and inference: a practical information-theoretic approach (2^nd^ edition) New York, NY: Springer.

Cassaing J., B. Le Proux de la Riviere, F. De Donno, E. Martinez-Garcia, and C. Thomas. 2013. Interactions between 2 mediterranean rodent species: habitat overlap and use of heterospecific cues. Écoscience 20:137–147.

Cayuela, H., O. Grolet, and P. Joly. 2018. Context-dependent dispersal, public information and heterospecific attraction in newts. Oecologia 188:1069–1080.

Chaianunporn, T. and T. Hovestadt. 2012. Evolution of dispersal in metacommunities of interacting species. Journal of Evolutionary Biology 24:2511–2525.

Clobert, J., J. F. Le Galliard, J. Cote, S. Meylan, and M. Massot. 2009. Informed dispersal, heterogeneity in animal dispersal syndromes and the dynamics of spatially structured populations. Ecology Letters 12:197–209.

Danchin, É., L. A. Giraldeau, T. J. Valone, and R. H. Wagner. 2004. Public information: from nosy neighbors to cultural evolution. Science 305:487–491.

Danielson, B.J. and M.S. Gaines, 1987. The influences of conspecific and heterospecific residents on colonization. Ecology 68:1778–1784.

De Bona, S., M. Bruneaux, A. E. G. Lee, D. N. Reznick, P. Bentzen, and A. López-Sepulcre. 2019. Spatio-temporal dynamics of density-dependent dispersal during a population colonisation. Ecology Letters 22:634–644.

De Meester, N., S. Derycke, A. Rigaux, and T. Moens. 2015. Active dispersal is differentially affected by inter- and intraspecific competition in closely related nematode species. Oikos 124:561–570.

Denomme-Brown, S.T., K. Cottenie, J. B. Falls, E. A. Falls, R. J. Brooks, and A. G. McAdam. Accepted to Oecologia pending revisions. Variation in space and time: a long-term examination of density-dependent dispersal in a woodland rodent.

Do, R., J. Shonfield, and A. G. McAdam. 2013. Reducing accidental shrew mortality associated with small-mammal livetrapping II: a field experiment with bait supplementation. Journal of Mammalogy 94:754–760.

Doligez, B., T. Pärt, E. Danchin, J. Clobert, and L. Gustafsson. 2004. Availability and use of public information and conspecific density for settlement decisions in the collared flycatcher. Journal of Animal Ecology 73:75–87.

Ford, A.T. and L. Fahrig. 2008. Movement patterns of eastern chipmunks (*Tamias striatus*) near roads. Journal of Mammalogy 89:895–903.

Gaines, M.S. and L.R. McClenaghan. 1980. Dispersal in small mammals. Annual Review of Ecology, Evolution and Systematics 11:163–196.

Getty, T. 1981. Structure and dynamics of chipmunk home range. Journal of Mammalogy 62:726–737.

Greenwood, P.J. 1980. Mating systems philopatry and dispersal in birds and mammals. Animal Behavior 28:1140–1162.

Hauzy, C., F. D. Hulot, A. Gins, and M. Loreau. 2007. Intra- and interspecific density-dependent disperal in an aquatic prey-predator system. Journal of Animal Ecology 76:552–558.

Heeb, P. et al. 1999. Ectoparasite infestation and sex-biased local recruitment of hosts. Nature 400:63–65.

Hestbeck, J. B. 1982. Population regulation of cyclic mammals: the social fence hypothesis. Oikos 39:157–163.

Holekamp, K. E. 1986. Proximal causes of natal dispersal in Belding’s ground squirrels (*Spermophilus beldingi*). Ecological Monographs 56:365–391.

Holyoak, M., R. Casagrandi, R. Nathan, E. Revilla, and O. Spiegel. 2008. Trends and missing parts in the study of movement ecology. Proceedings of the National Academy of Sciences 105: 19060–19065.

Howard, W.E. Innate and environmental dispersal of individual vertebrates. The American Midland Naturalist 63(1):152–161.

Koenig, W., D. Van Vuren, and P. N. Hooge. 1996. Detectability, philopatry and the distribution of dispersal distances in vertebrates. Trends in Ecology & Evolution 11:514–517.

Lacher, T.E. and M.A. Mares. 1996. Availability of resources and use of space in eastern chipmunks, *Tamias striatus*. Journal of Mammalogy 77:833–849.

Matthysen, E. 2005. Density-dependent dispersal in birds and mammals. Ecography 28:403–416.

Mönkkönen, M., R. Härdling, J. T. Forsman, and J. Tuomi. 1999. Evolution of heterospecific attraction: using other species as cues in habitat selection. Ecology 13:91–104.

Nathan, R. 2006. Long-distance dispersal of plants. Science 313:786–788.

Ovaska, K. and T.B. Herman. 1988. Life history characteristics and movements of the woodland jumping mouse. *Napaeozapus insignis*, in Nova Scotia. The Canadian Journal of Zoology 66:1752–1762.

Parejo, D., É. Danchin, and J. M. Avilés. 2004. The heterospecific habitat copying hypothesis: can competitors indicate habitat quality? Behavioral Ecology 16:96–105.

Parejo, D., É. Danchin, N. Silva, J. F. White, N. Amélie, and J. M. Avilés. 2008. Do great tits rely on inadvertent social information from blue tits? A habitat selection experiment. Behavioral Ecology and Sociobiology 62:1569–1579.

Porter, J.H. and J.L. Dooley. 1993. Animal dispersal patterns: a reassessment of simple mathematical models. Ecology 74:2436–2443.

Rehmeier, R. L., G. A. Kaufman, and D. W. Kaufman. 2004. Long-distance movements of the deer mouse in tallgrass prairie. Journal of Mammalogy 85:562–568.

Seppänen, J.T., J. T. Forsman, M. Mönkkönen, and R. L. Thomson. 2007. Social information use is a process across time, space, and ecology, reaching heterospecifics. Ecology 88:1622–1633.

Shonfield, J., R. Do, R. J. Brooks, and A. G. McAdam. 2013. Reducing accidental shrew mortality associated with small-mammal livetrapping I: an inter- and intrastudy analysis. Journal of Mammalogy 94:745–753.

Sikes, R.S. 2016. 2016 Guidelines of the American Society of Mammalogists for the use of wild mammals in research and education. Journal of Mammalogy 97:663–688.

Sloggett J.J. and W.W. Weisser. 2002. Parasitoids induce production of the dispersal morph of the pea aphid, *Acyrthosiphon pisum*. Oikos 98:323–333.

Stamps, J. A. 1991. The effects of conspecifics on habitat selection in teritorial species. Behavioral Eoclogy and Sociobiology 28:29–36.

Sutherland, G. D., A. S. Harestad, K. Price, and K. P. Lertzman. 2000. Scaling of natal dispersal distances in terrestrial birds and mammals. Conservation Ecology 4: 16.

Sutherland, W. J. et al. 2013. Identification of 100 fundamental ecological questions. Journal of Ecology 101:58–67.

Szymkowiak, J., R. L. Thomson, and L. Kuczyński. 2017. Interspecific social information use in habitat selection decisions among migrant songbirds. Behavioral Ecology 28:767–775.

Tilman, D. 1987. The importance of the mechanisms of interspecific competition. The American Naturalist 129:769–774.

Valone, T.J. 1989. Group foraging, public information and patch estimation. Oikos 56:357–363.

Valone T.J. and J.J. Templeton. 2002. Public information for the assessment of quality: a widespread social phenomenon. Philosophical Transactions of the Royal Society B: Biological Sciences 357:1549–1557.

van de Pol, M. and J. Wright. 2009. A simple method for distinguishing within-versus between-subject effects using mixed models. Animal Behaviour 77: 753–758.

Vignieri, S.N. 2005. Streams over mountains: influence of riparian connectivity on gene flow in the pacific jumping mouse (*Zapus trinotatus*). Molecular Ecology 14:1925–1937.

Vignieri, S.N. 2007. Cryptic behaviours, inverse genetic landscapes and spatial avoidance of inbreeding in the pacific jumping mouse. Molecular Ecology 16:853–866.

