## Supplemental Materials 1-5 for "Examining Interspecific density-dependent dispersal in forest small mammals"

### SUPPLEMENTARY DATA

*SD1—conspecific density dependent dispersal candidate model structure*

Below is listed each candidate conspecific density-dependent dispersal model for each species in this manuscript and their corresponding fixed effects. The structure of the candidate models was consistent for each species. All models used whether an individual of the species of study dispersed or did not as the response variable.

Candidate Model Fixed Effects:

1) Distance to Closest Line (km)
2) Distance to Closest Line (km), Sex (M/F)
3) Distance to Closest Line (km), Sex (M/F), Local Density
4) Distance to Closest Line (km), Sex (M/F), Regional Density,
5) Distance to Closest Line (km), Sex (M/F), Relative Local Density
6) Distance to Closest Line (km), Sex (M/F), Relative Local Density, Regional Density
7) Distance to Closest Line (km), Sex (M/F), Relative Local Density:Regional Density
8) Distance to Closest Line (km), Sex (M/F), Relative Local Density:Regional Density, Relative Local Density:Sex(M/F)

In the case of red-backed vole, in model 3) a quadratic term rather than a linear term was fitted to account for a significant nonlinearity.

*SD2—conspecific density dependent dispersal model AIC comparisons*

Below are tables containing AIC comparisons for conspecific density-dependent dispersal models for each species examined in this study. Model numbers refer to the structure laid out in SD1.

Table 1: Eight generalized linear mixed effects models (binomial response, logit link) fitted to data for predicting the probability of individual red-backed vole dispersal based on conspecific density

| Model | *K* | *AIC* | *ΔAIC* | *Rank* |
|---|---|---|---|---|
| 1) | 4 | 198.6 | 0 | 1 |
| 2) | 5 | 199.3 | 0.7 | 2 |
| 3) | 7 | 199.5 | 0.9 | T-3 |

| | | | | |
|---|---|---|---|---|
| 7) | 8 | 199.5 | 0.9 | T-3 |
| 5) | 6 | 200.6 | 2.0 | 5 |
| 4) | 6 | 201.2 | 2.6 | 6 |
| 8) | 9 | 201.5 | 2.9 | 7 |
| 6) | 7 | 202.5 | 3.9 | 8 |

K = number of parameters, AIC = Akaike Information Criterion, ΔAIC = Model's AIC - the lowest AIC of any model

Random effects in both models were *year* and *trapline of emigration*.

Table 2: Eight generalized linear mixed effects models (binomial response, logit link) fitted to data for predicting the probability of individual eastern chipmunk dispersal based on conspecific density

| Model | *K* | *AIC* | *ΔAIC* | *Rank* |
|---|---|---|---|---|
| 2) | 5 | 329.5 | 0 | 1 |
| 1) | 4 | 329.9 | 0.4 | 2 |
| 4) | 6 | 330.5 | 1.0 | 3 |
| 5) | 6 | 330.7 | 1.2 | 4 |
| 3) | 6 | 331.2 | 1.7 | 5 |
| 6) | 7 | 332.7 | 3.2 | 6 |
| 8) | 9 | 334.0 | 4.5 | 7 |
| 7) | 8 | 334.6 | 5.1 | 8 |

K = number of parameters, AIC = Akaike Information Criterion, ΔAIC = Model's AIC - the lowest AIC of any model

Random effects in both models were *year* and *trapline of emigration*.

Table 3: Eight generalized linear mixed effects models (binomial response, logit link) fitted to data for predicting the probability of individual jumping mice dispersal based on conspecific density

| Model | *K* | *AIC* | *ΔAIC* | *Rank* |
|---|---|---|---|---|
| 3) | 6 | 212.0 | 0 | 1 |
| 5) | 6 | 212.4 | 0.4 | 2 |
| 6) | 7 | 213.8 | 1.8 | 3 |

| | | | | |
|---|---|---|---|---|
| 7) | 8 | 215.5 | 3.5 | 4 |
| 8) | 9 | 217.4 | 5.4 | 5 |
| 2) | 5 | 217.9 | 5.9 | 6 |
| 4) | 9 | 218.8 | 6.8 | 7 |
| 1) | 4 | 220.5 | 8.5 | 8 |

K = number of parameters, AIC = Akaike Information Criterion, ΔAIC = Model's AIC - the lowest AIC of any model

Random effects in both models were *year* and *trapline of emigration*.

*SD3—heterospecific density dependent dispersal candidate model structure and output*

Below are listed the candidate heterospecific density-dependent dispersal models for each species in this study and their corresponding fixed effects. For each species the "base model" refers to the best model from the conspecific analysis for each species. All models used whether an individual of the species of study dispersed or did not as the response variable.

Fixed effects in candidate heterospecific density-dependent dispersal models for deer mice:

1) Base Model Effects
2) Base Model Effects, Local Density of eastern chipmunk
3) Base Model, Local Density of red-backed vole
4) Base Model, Local Density of jumping mice
5) Base Model, Cumulative local density of other three species (local competitor density)
6) Base Model, local competitor density, local competitor density:Sex(M/F)

Fixed Effects in candidate heterospecific density-dependent dispersal models for eastern chipmunk:

1) Base Model Effects
2) Base Model Effects, Local Density of deer mice
3) Base Model, Local Density of red-backed vole
4) Base Model, Local Density of jumping mice
5) Base Model, Cumulative local density of other three species (local competitor density)
6) Base Model, Local competitor density,Local competitor density:Sex(M/F)

Fixed Effects in candidate heterospecific density-dependent dispersal models for red-backed vole:

1) Base Model Effects

2) Base Model Effects, Local Density of deer mice
3) Base Model, Local Density of eastern chipmunk
4) Base Model, Local Density of jumping mice
5) Base Model, Cumulative local density of other three species (local competitor density)
6) Base Model, Local competitor density, Local competitor density:Sex(M/F)

Fixed Effects in candidate heterospecific density-dependent dispersal models for jumping mice:

1) Base Model Effects
2) Base Model Effects, Local Density of deer mice
3) Base Model, Local Density of eastern chipmunk
4) Base Model, Local Density of red-backed vole
5) Base Model, Cumulative local density of other three species (local competitor density)
6) Base Model, Local competitor density, Local competitor density:Sex(M/F)

*Heterospecific density dependent dispersal model AIC comparisons*

Below are tables containing AIC comparisons for each conspecific density-dependent dispersal model for each species examined in this study. Model numbers refer to the structure in the previous section I.d).

Table 1: Seven generalized linear mixed effects models (binomial response, logit link) fitted to data for predicting the probability of individual deer mice dispersal based on heterospecific density

| Model | *K* | *AIC* | *ΔAIC* | *Rank* |
|---|---|---|---|---|
| 1) | 9 | 1059.6 | 0 | 1 |
| 3) | 10 | 1061.3 | 1.7 | 2 |
| 5) | 10 | 1061.5 | 1.9 | 3 |
| 2) | 10 | 1061.6 | 2.0 | T-4 |
| 4) | 10 | 1061.6 | 2.0 | T-4 |
| 6) | 11 | 1062.6 | 3.0 | 6 |

K = number of parameters, AIC = Akaike Information Criterion, ΔAIC = Model's AIC - the lowest AIC of any model

Random effects in both models were *year* and *trapline of emigration*.

Table 2: Seven generalized linear mixed effects models (binomial response, logit link) fitted to data for predicting the probability of individual eastern chipmunk dispersal based on heterospecific density

| Model | *K* | *AIC* | *ΔAIC* | *Rank* |
|---|---|---|---|---|
| 1) | 6 | 327.1 | 0 | 1 |
| 2) | 7 | 328.2 | 1.1 | 2 |
| 5) | 7 | 328.3 | 1.2 | T-3 |
| 4) | 7 | 328.3 | 1.2 | T-3 |
| 3) | 7 | 328.8 | 1.7 | 5 |
| 6) | 8 | 330.1 | 3.0 | 6 |

K = number of parameters, AIC = Akaike Information Criterion, ΔAIC = Model's AIC - the lowest AIC of any model

Random effects in both models were *year* and *trapline of emigration*.

Table 3: Seven generalized linear mixed effects models (binomial response, logit link) fitted to data for predicting the probability of individual red-backed vole dispersal based on heterospecific density

| Model | *K* | *AIC* | *ΔAIC* | *Rank* |
|---|---|---|---|---|
| 1) | 8 | 198.6 | 0 | 1 |
| 4) | 9 | 198.7 | 0.1 | 2 |
| 3) | 9 | 199.8 | 1.2 | 3 |
| 2) | 9 | 200.6 | 2.0 | T-4 |
| 5) | 9 | 200.6 | 2.0 | T-4 |
| 6) | 10 | 202.0 | 3.4 | 6 |

K = number of parameters, AIC = Akaike Information Criterion, ΔAIC = Model's AIC - the lowest AIC of any model

Random effects in both models were *year* and *trapline of emigration*.

Table 4: Seven generalized linear mixed effects models (binomial response, logit link) fitted to data for predicting the probability of individual jumping mice dispersal based on heterospecific density

| Model | *K* | *AIC* | *ΔAIC* | *Rank* |
|---|---|---|---|---|
| 3) | 7 | 210.2 | 0 | 1 |
| 6) | 8 | 211.4 | 1.2 | 2 |
| 5) | 7 | 211.6 | 1.4 | 3 |
| 1) | 6 | 212.0 | 1.8 | 4 |
| 4) | 7 | 213.1 | 2.9 | 5 |
| 2) | 8 | 213.8 | 3.6 | 6 |

K = number of parameters, AIC = Akaike Information Criterion, ΔAIC = Model's AIC - the lowest AIC of any model
Random effects in both models were *year* and *trapline of emigration*.

*SD4—summary statistics regarding deer mice in APP (taken from Denomme-Brown et al. in review)*

Denomme-Brown et al. (*in review*) reported 3408 individual deer mice caught a minimum of 3 times, hereafter termed possible dispersing individuals. Over their study 4.2% (n=142) of possible dispersers were detected dispersing. Distances travelled ranged from 115m to 11.4km ($\bar{x} \pm SE = 1.68 \pm 0.20$ km).

Density of deer mice varied over an order of magnitude throughout the study. Annual regional population densities ranged between 1.9 and 27.8 captures per hundred trapnights ($\bar{x} \pm SE = 11.15 \pm 0.81$ captures per hundred trap-nights, n = 51 years). Local density within years was also highly variable ($\bar{x} \pm SE = 19.37 \pm 1.22$ captures per hundred trap-nights, n = 51 years).

*SD5—Full model results for consistently structured heterospecific density-dependent dispersal models*

Table 1: Summary of the statistical results from deer mice heterospecific density-dependent dispersal model from consistently structured model comparison. This model best fitted the observed dispersal data and had strong support based on model comparisons using AIC. Significant effects (P < 0.05) are in boldface. Likelihood ratio tests were used to assess the significance of each random effect.

| *Fixed* | *β ± SE* | *Z* | *P* |
|---|---|---|---|
| Closest Distance | -0.99 ± 0.17 | -5.98 | **<0.0001** |
| Local Conspecific Density | -0.50 ± 0.13 | -3.89 | **0.0001** |
| Local Chipmunk Density | -0.099 ± 0.15 | -0.68 | 0.49 |

| | | | |
|---|---|---|---|
| Local Vole Density | 0.045 ± 0.084 | 0.53 | 0.60 |
| Local Jumping Mice Density | 0.052 ± 0.10 | 0.54 | 0.60 |

| *Random* | $\partial^2$ | $X^2{}_1$ | *P* |
|---|---|---|---|
| Year | 1059 | $11.1_1$ | **0.0009** |
| Initial Trapline | 1070 | $0.1_1$ | 0.75 |

Table 2: Summary of the statistical results from eastern chipmunk heterospecific density-dependent dispersal model from consistently structured model comparison. This model best fitted the observed dispersal data and had strong support based on model comparisons using AIC. Significant effects ($P < 0.05$) are in boldface. Likelihood ratio tests were used to assess the significance of each random effect.

| *Fixed* | $\beta \pm SE$ | *Z* | *P* |
|---|---|---|---|
| Closest Distance | -1.37 ± 0.27 | -5.03 | **<0.0001** |
| Local Conspecific Density | -0.029 ± 0.23 | -0.13 | 0.90 |
| Local Deer Mice Density | -0.14 ± 0.18 | -0.78 | 0.43 |
| Local Vole Density | 0.015 ± 0.15 | -0.10 | 0.92 |
| Local Jumping Mice Density | -0.18 ± 0.18 | -0.98 | 0.33 |

| *Random* | $\partial^2$ | $X^2{}_1$ | *P* |
|---|---|---|---|
| Year | 318 | $0_1$ | 1 |
| Initial Trapline | 318 | $0_1$ | 1 |

Table 3: Summary of the statistical results from red-backed vole heterospecific density-dependent dispersal model from consistently structured model comparison. This model best fitted the observed dispersal data and had strong support based on model comparisons using AIC. Significant effects ($P < 0.05$) are in boldface. Likelihood ratio tests were used to assess the significance of each random effect.

| *Fixed* | *β ± SE* | *Z* | *P* |
|---|---|---|---|
| Closest Distance | -0.33 ± 0.56 | -0.58 | 0.56 |
| Local Conspecific Density | -0.16 ± 0.29 | -0.55 | 0.59 |
| Local Deer Mice Density | -0.11 ± 0.28 | -0.40 | 0.69 |
| Local Eastern Chipmunk Density | 0.32 ± 0.24 | 1.34 | 0.18 |
| Local Jumping Mice Density | -0.47 ± 0.32 | -1.45 | 0.15 |

| *Random* | $\partial^2$ | $X^2_1$ | *P* |
|---|---|---|---|
| Year | 188 | $0.15_1$ | 0.70 |
| Initial Trapline | 187 | $0.87_1$ | 0.35 |

Table 4: Summary of the statistical results from jumping mice heterospecific density-dependent dispersal model from consistently structured model comparison. This model best fitted the observed dispersal data and had strong support based on model comparisons using AIC. Significant effects ($P < 0.05$) are in boldface. Likelihood ratio tests were used to assess the significance of each random effect.

| *Fixed* | *β ± SE* | *Z* | *P* |
|---|---|---|---|
| Closest Distance | -0.66 ± 0.38 | -1.76 | 0.08 |
| Local Conspecific Density | -0.62 ± 0.27 | -2.32 | **0.02** |
| Local Deer Mice Density | -0.12 ± 0.25 | 0.49 | 0.62 |
| Local Eastern Chipmunk Density | -0.94 ± 0.49 | -1.93 | 0.05 |
| Local Vole Density | -0.17 ± 0.23 | -0.75 | 0.45 |

| *Random* | $\partial^2$ | $X^2_1$ | *P* |
|---|---|---|---|

| | | | |
|---|---|---|---|
| Year | 202 | $0_1$ | 1 |
| Initial Trapline | 201 | $1.21_1$ | 0.27 |
